# Choosing priors in Bayesian ecological models by simulating from the prior predictive distribution

**DOI:** 10.1101/2020.12.10.419713

**Authors:** Jeff S. Wesner, Justin P.F. Pomeranz

## Abstract

Bayesian data analysis is increasingly used in ecology, but prior specification remains focused on choosing non-informative priors (e.g., flat or vague priors). One barrier to choosing more informative priors is that priors must be specified on model parameters (e.g., intercepts, slopes, sigmas), but prior knowledge often exists on the level of the response variable. This is particularly true for common models in ecology, like generalized linear mixed models, which may have a link function and dozens of parameters, each of which needs a prior distribution. We suggest that this difficulty can be overcome by simulating from the prior predictive distribution and visualizing the results on the scale of the response variable. In doing so, some common choices for non-informative priors on parameters can easily be seen to produce biologically impossible values of response variables. Such implications of prior choices are difficult to foresee without visualization. We demonstrate a workflow for prior selection using simulation and visualization with two ecological examples (predator-prey body sizes and spider responses to food competition). This approach is not new, but its adoption by ecologists will help to better incorporate prior information in ecological models, thereby maximizing one of the benefits of Bayesian data analysis.

## Introduction

The distinguishing feature between Bayesian and non-Bayesian statistics is that Bayesian statistics treats unknown parameters as random variables governed by a probability distribution, while non-Bayesian statistics treats unknown parameters as fixed (Ellison and Dennis 2010, Hobbs and Hooten 2015). A common misconception is that only Bayesian statistics incorporates prior information. However, non-Bayesian methods can and often do incorporate prior information, either informally in the choices of likelihoods and model structures, or formally as penalized likelihood or hierarchical modeling (Hobbs and Hooten 2015, Morris et al. 2015).

While prior information is not unique to Bayesian models, it is required of them. For example, in a simple linear regression of the form *y* ~ *N* (*α* + *βx*, *σ*), the intercept *α*, slope *β*, and error *σ* are unknown parameters that need a prior probability distribution. There are differing opinions and philosophies on the best practices for choosing priors (Lindley 1961, Edwards et al. 1963, Morris et al. 2015, Wolf et al. 2017, Lemoine 2019, Banner et al. 2020, Gelman et al. 2017). In ecology, a common practice is to assign so-called non-informative priors that effectively assign equal probability to all possible values using either uniform or diffuse normal priors with large variances (Lemoine 2019). These priors allow Bayesian inference to proceed (i.e. produce a posterior distribution), but with presumably limited influence of the priors (Gelman et al. 2013, Lemoine 2019).

Reasons for using non-informative priors are varied but are at least in part driven by a desire to avoid the appearance of subjectivity and/or a reliance on default settings in popular software (Gelman and Hennig 2017, Banner et al. 2020). There are several current arguments against this approach. First, “non-informative” is a misnomer. All priors influence the posterior distribution to some extent (Hobbs and Hooten 2015). As a result, a prior cannot just be assumed as non-informative based on default settings or a wide variance (Seaman III et al. 2012). Its implications for the model should be checked just like any other subjective assumption in data analysis, whether Bayesian or not (Banner et al. 2020, Gelman et al. 2017). Second, adhering to non-informative priors removes a major potential benefit of Bayesian analysis, which is to explicitly incorporate prior research and expertise into new science (Hobbs and Hooten 2015, Lemoine 2019, Rodhouse et al. 2019). Third, informative priors can help to reduce spurious conclusions due to errors in magnitude or sign of an effect by treating extreme values in the data skeptically (Gelman et al. 2012, Lemoine 2019). Finally, informative priors make computational algorithms like MCMC run more efficiently, which can save hours or days of computing time in complex models (Hobbs and Hooten 2015).

However, while there are clear arguments for why ecologists *should* use more informative priors, it is often difficult to know *how* to use them. Even for seemingly simple and routine models, like logistic regression, it can be difficult to understand *a priori* how priors affect the model, because they must be assigned in the context of likelihood with a linearizing link-function (Seaman III et al. 2012, Gelman et al. 2017). In other words, prior specification takes place on model parameters (e.g., slopes, intercepts, variances), but prior knowledge is often easier to assess on the model outcomes (Kadane et al. 1980, Bedrick et al. 1996, Gabry et al. 2019). This is particularly true for the types of models that are commonly used in ecology, such as generalized linear mixed models with interactions, which may have dozens of parameters and hyperparameters, each of which require a prior probability distribution (Bedrick et al. 1996, McElreath 2020).

We suggest that ecologists can address this problem using simulation from the prior predictive distribution and visualizing the implications of the priors on either the expected mean of the data (e.g., simulate regression lines or group means) (Kadane et al. 1980, Bedrick et al. 1996) or by simulating individual data points (Gabry et al. 2019). In this paper, we demonstrate this approach using two case studies with ecological data. All data and code for these examples, as well as additional case studies, are available at: https://github.com/jswesner/prior_predictive.

### Prior Predictive Simulation

An attractive feature of the Bayesian approach is that the models are generative. This means that we can simulate potential data from the model so long as the parameters are assigned a proper probability distribution (Gelman et al. 2013). This feature is routinely used to check models and prior influence *after* fitting the data using the posterior predictive distribution (Lemoine 2019, Gelman et al. 2020), but it can also be used before seeing the data using the prior predictive distribution (Gabry et al. 2019). As an example, consider a simple linear regression:

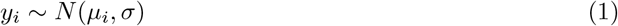

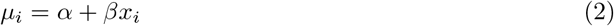

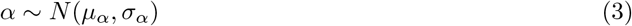

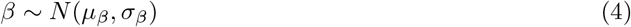

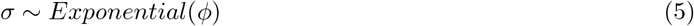

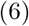

where *y_i_* is the response variable on the *i^th^* row, normally distributed with an unknown mean *µ_i_* and standard deviation *σ*. We assume that the response variable is a linear function of the predictor variable *x_i_* using a linear equation with an intercept *α* and a slope *β*. The following parameters need priors: *σ, α, β*. The first step is to choose a probability distribution for each prior. We chose normal (i.e. Gaussian) distributions for *α* and *β* and an exponential distribution for *σ*. The normal distributions imply that the intercept or slope are continuous and can be positive or negative (Hobbs and Hooten 2015). The exponential distribution is a common prior for standard deviations because it generates only positive values and allows for occasionally large deviations. For standard deviations (or variances or precisions), there are a number of alternatives prior distributions available (Gelman and others 2006, Gelman et al. 2013, McElreath 2020).

The challenge is to assign prior values to the mean and sd of each normal distribution and to *ϕ* of the exponential distribution before seeing the data *y_i_*. One way to do that is to use prior parameter estimates from previous studies (Rodhouse et al. 2019). For example, if previous studies found that the slope *β* was typically 1.2 with a standard deviation of 0.5 for *σ_β_*, then we could use *β* ~ *N* (1.2, 0.5) as the prior for this parameter. Similarly, if other studies suggested that the residuals were described by an exponential distribution with a rate parameter *ϕ* of 0.34, then we could use that here.

In our experience it is more common that the current study might differ slightly from previous studies, either in the experimental design, the spatial scale, the species used, or the inclusion of additional covariates. Those differences make it more difficult to use a posterior distribution of a parameter from one model as a prior distribution in another model. Simulation offers a practical approach here.

The general workflow for prior predictive simulation is:

1. Draw N values from different prior distributions. (e.g., *α_sim_ N* (0, 100)), *β_sim_ N* (0, 10)…, *σ exp*(0.1))
2. For each draw, solve the equation for each *i* value of *x*. (e.g., *µ_i_* = *α* + *βx_i_*))
3. Plot the result for either *µ_i_*, *y_i_*, or another derived quantity. (e.g., 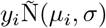)
4. Use our domain knowledge (or another expert’s) to assess whether the simulated values reflect prior knowledge.
5. If simulated values do not reflect prior knowledge, change the prior distribution, likelihood, or both and repeat the simulation from step 1.
6. If simulated values reflect prior knowledge, add the data and estimate the posterior distribution.

This amounts to a prior predictive check to satisfy the expectation that “simulations from the full Bayesian model. ‥ should be plausible data sets” (Kennedy et al. 2019). The simulation and visualization steps (1-3) are critical, because simulated data sets are derived from the *joint distribution* of parameters. In other words, whether a model simulates plausible data data cannot be determined simply from looking at the individual priors or model formula, because their interpretation depends on the units of measurement (e.g., a N(0,1) prior means different things if *y* is measure in *µ*m versus km) and on the range of prior expected values. For step 5, it is important to emphasize that there is no precise definition for what “ reflects prior knowledge”. The purpose of prior simulation is not to pre-determine an outcome, but instead to make explicit exactly how and why the priors were chosen. The importance of those priors on posterior inference should still be assessed, but that topic is beyond the scope of this paper. We demonstrate prior predictive simulation below with two motivating examples.

## Motivating Examples

### Example 1: Predator-Prey Body Sizes - Simple Linear Regression

#### Data

Understanding predator-prey interactions has long been a research interest of ecologists. Body size is related to a number of aspects that influence these interactions. For example, predators are often gape-limited, meaning that larger predators should be able to eat larger prey. The data set of (Brose et al. 2006) documents over 10,000 predator-prey interactions, including the mean mass of each.

#### Model

For this example, we examine the hypothesis that the mean prey body mass increases log-linearly with predator body mass using a simple linear model:

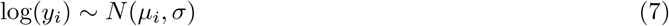

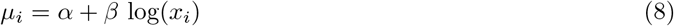

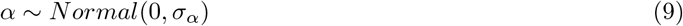

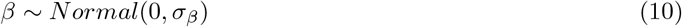

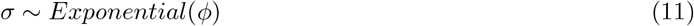

where log(*y_i_*) is natural log transformed prey mass and log(*x_i_*) is natural log transformed predator mass.

#### Priors

For the *α* and *β* priors, we need to specify a mean and standard deviation. As a first guess, we assign a mean of 0 with a “non-informative” standard deviation of 1000 [*N* (0, 1000)]. The mean of 0 in a normal distribution implies that the intercept and slope have equal probability of being positive or negative. For the standard deviation, a rule of thumb is to assume that reasonable values can be anywhere between *±* twice the standard deviation (McElreath 2020). There is nothing special about this prior *per se*, but it was a common default setting in earlier Bayesian software to generate “flat” prior distributions (usually specified as a precision rather than a standard deviation) and appears regularly in the literature (McCarthy and Masters 2005, Banner et al. 2020). Similarly, for the exponential distribution, smaller rates *ϕ* generate larger deviations, so we’ll specify an initial *ϕ* of 0.00001. We chose this initial value by plotting 100 simulations from the exponential function in R (R Core Team 2020) under varying values of *ϕ* [e.g., `plot(rexp(100, 0.00001)`]. A value of 0.00001 generated an average deviance of ~1,000 with values up to ~5,000, indicating the possibility of producing extremely large values.

After simulating from these initial priors, we specified successfully tighter priors (Table 1). We then simulated from the prior predictive distribution and compared those simulations to reference points representing prior knowledge (Mass of earth, a Blue Whale, a virus, and a Carbon-12 atom). The goal was to use these reference points to find a joint prior distribution that produced reasonable values of potential prey masses. We did this using two levels of the model (*µ_i_* and *y_i_*). For *µ_i_*, we simulated 100 means across each value of *x_i_* and plotted them as regression lines. For *y_i_*, we simulated a fake data set containing simulated values of log prey mass for each of the 13,085 values of log predator mass (*x_i_*) in the (Brose et al. 2006) data.

**Table 1:** Priors used for the two models. Distributions are either normal with a mean and standard deviation [N(mu, sigma)] or exponential [Exp(rate)].

| Parameter | Model 1: Predator-Prey |  |  | Model 2: Spiders |  |  |
| --- | --- | --- | --- | --- | --- | --- |
|  | Weak | Strong | Strongest | Weak | Strong | Strongest |
| Alpha | $N(0,1000)$ | $N(0,10)$ | $N(0,1)$ | $N(0,10)$ | $N(0,1)$ | $N(0,0.1)$ |
| Beta(s) | $N(0,1000)$ | $N(0,10)$ | $N(0,1)$ | $N(0,10)$ | $N(0,1)$ | $N(0,0.1)$ |
| Sigma | $Exp(0.001)$ | $Exp(0.01)$ | $Exp(0.1)$ | | | |
| Sigma_alpha | | | | $Exp(0.1)$ | $Exp(1)$ | $Exp(2)$ |
| Sigma_cage | | | | $Exp(0.1)$ | $Exp(1)$ | $Exp(2)$ |

#### Results

Based on simulations from the prior predictive distribution, the weak “non-informative” priors make nonsense predictions (Figure 1a-c). In Figure 1a, all of the lines are impossibly steep, suggesting that predators could plausibly eat prey that are much larger than earth or much smaller than an atom. The seemingly stronger priors in Figure 1b suffer from the same problem, though the effect is less severe. The strongest priors (Figure 1c) produce more reasonable predictions, though they are still quite vague, with positive probability that large and small predators could eat prey that are orders of magnitude larger than an adult Blue Whale. The simulated fake data sets tell a similar story (Figure 1d-f), but with the added influence of *sigma* in the likelihood (Equation 6).

**Figure 1:**
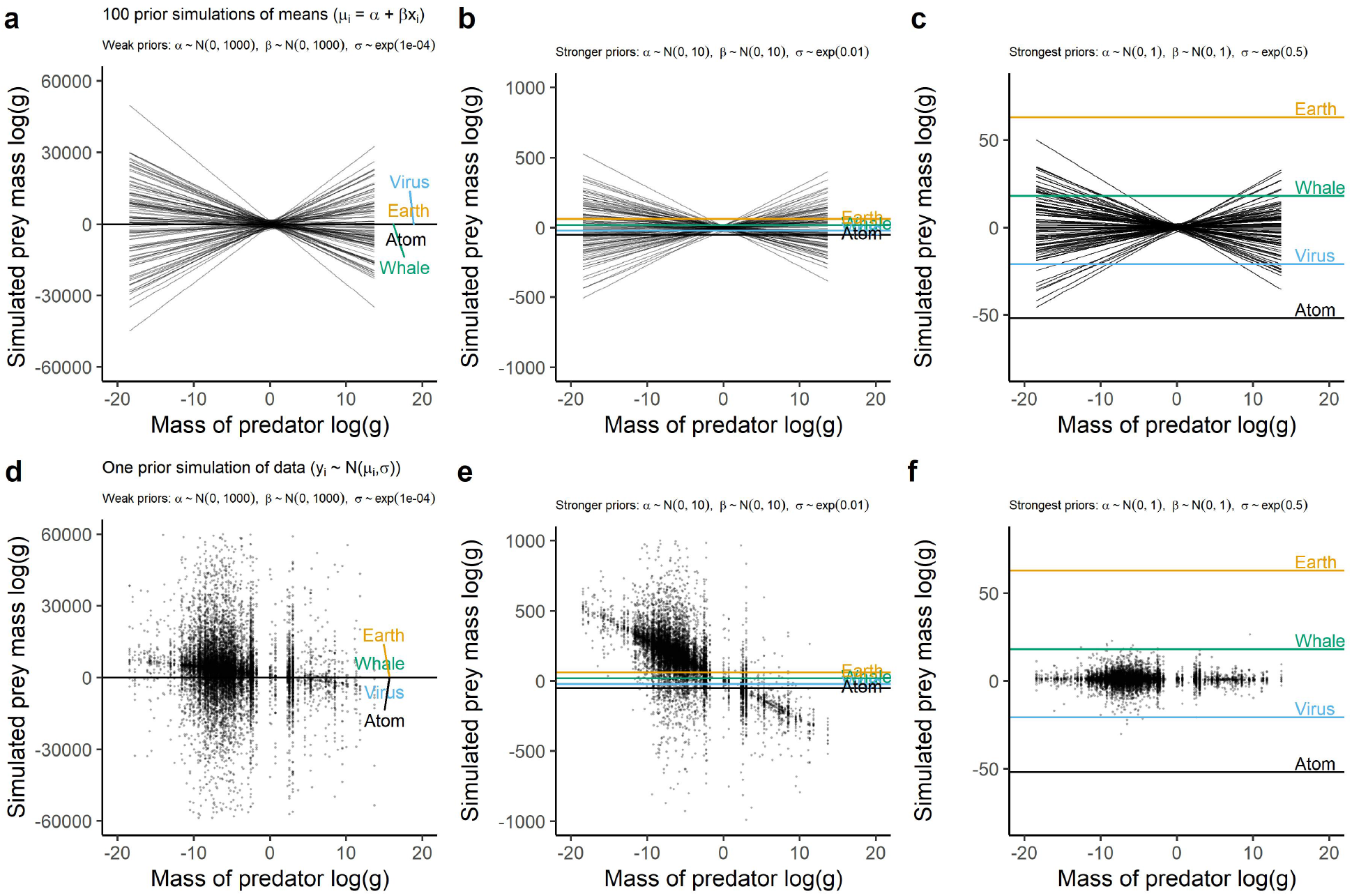
Prior predictive simulations showing the implications of the priors on predictions of log prey mass. The top row (a-c) shows prior simulations of regression lines. The bottom row (d-f) shows prior predictive simulation of one dataset.

We fit the model using the strongest prior set (Figure 2). As is typical of models with large amounts of data, the parameters (i.e. slope, intercept, and sigma) are similar regardless of the prior distribution (Figure 2). The intercept is −4.8 ± 0.04 (mean ± sd), the slope is 0.6 ± 0.01, and sigma is 3.7 ± 0.02.

**Figure 2:**
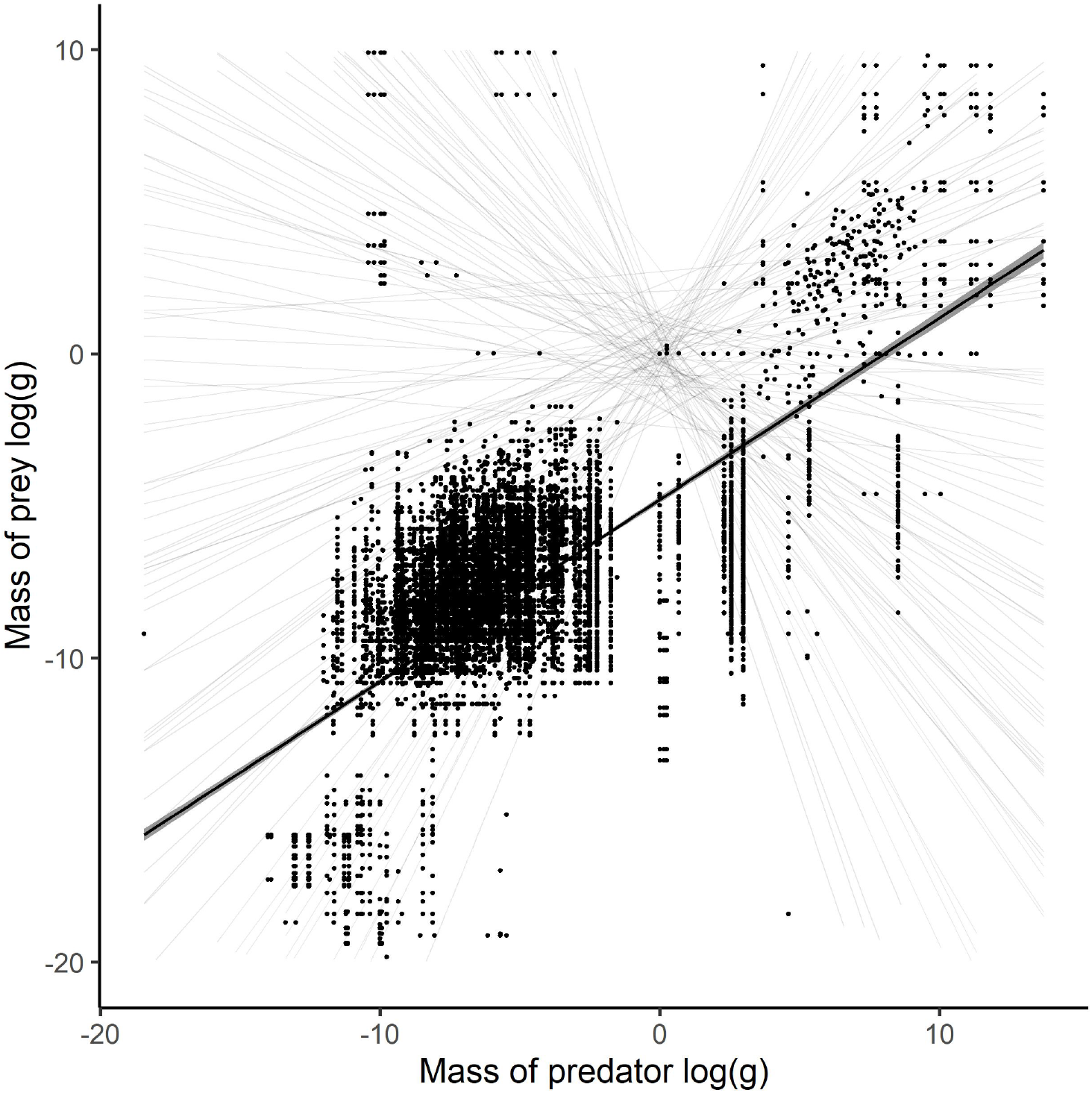
The fitted model showing the posterior distribution of the regression and raw data points from Brose et al. 2006. The shaded lines in the background are prior predictions from the strongest priors.

Regardless of the similarity in results, there are several benefits to choosing a stronger prior. First, it is difficult to justify the two weakest priors on biological grounds. They place large amounts of prior probability on impossible values. This can matter, for example, when priors need to be justified to a granting agency or to reviewers. More critically, specification of priors can have conservation or legal implications, and the ability to justify priors with simulation helps to improve transparency (Crome et al. 1996, Banner et al. 2020). Stronger priors also improve computational efficiency (McElreath 2020). We fit these models using the *brms* package (Burkner 2017). The models with stronger or strongest priors were up to 50% faster than the model with weak priors, taking 56 vs 28 seconds on a standard laptop (compilation time + warmup time + sampling time). For more complex models that take longer to run, this improvement can save hours or days of computing time.

#### Caveats

In a real analysis, there are some other steps we could have taken to generate a more realistic prior distribution before fitting the model to data. First, we know from the literature that predators are generally larger than their prey by 2-3 orders of magnitude (Trebilco et al. 2013). Therefore, it would make sense to alter the prior mean of the intercept to a value below zero, perhaps using an average predator/prey mass comparison from the literature. That is apparent from the prior versus posterior comparison in Figure 2. Similarly, the fact that larger predators tend to eat larger prey is well-known, so the prior on the slope *β* could be changed to a positive mean. One option would be to restrict the slope to only positive values, but this would not reflect our prior knowledge that predator body size is still a noisy predictor of prey body size (e.g., whales and parasitoids have prey that are orders of magnitude smaller or larger than they are, respectively).

Part of the uncertainty in prior selection can also be minimized by standardizing predictors (McElreath 2020). This changes the scale of each predictor so that the interpretation of its associated parameter is in units of standard deviation. In other words, a value of 2.3 for *β* would indicate that *y* increases by 2.3 for every standard deviation increase in *x*. Standardizing predictors can prevent problems that arise by mistaking cm for m or ha for acres. It also limits the expected prior values (e.g., N(0, 10) is extremely vague on a standardized predictor, but might be informative on a non-standardized predictor), but at the cost of less intuitive interpretation.

### Example 2: Spider Abundance - Generalized Linear Mixed Model

#### Data

This data set comes from (Warmbold and Wesner 2018), who studied how terrestrial spiders responded to different combinations of freshwater fish using fish enclosure cages in a backwater of the Missouri River, USA. The hypothesized mechanism was that fish would reduce the emergence of adult aquatic insects by eating the insects, causing a reduction in terrestrial spiders that feed on the adult forms of those insects. The original experiment contained six treatments. Here, we present a simplified version comparing spider abundance above three treatments that contain either Smallmouth Buffalo (*Ictiobus bubalus*), Green Sunfish (*Lepomis cyanellus*), or a fishless control. Each treatment had four replicates for a total of 12 cages (each 2.3 m^2^). The number of occupied spider webs above each cage were counted on four dates over the 29-day experiment.

#### Model

We fit a generalized linear mixed model with a Poisson likelihood, since the response variable (# webs) is a non-negative integer (i.e. number of spiders counted above a cage on each date). The predictor variables were date, treatment, and a date x treatment interaction. Since each replicate cage was sampled four times, we included a random intercept for cages. Describing the model as having two main effects and an interaction is deceptively simple. In reality, the model has 13 parameters that require a prior specification: 11 “fixed” effects that indicate all combinations of date x treatment, plus 1 intercept and a hyperprior *ϕ* on the intercept:

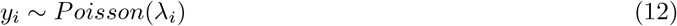

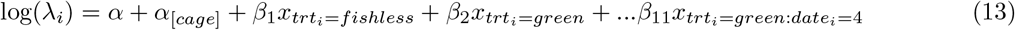

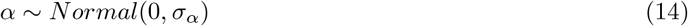

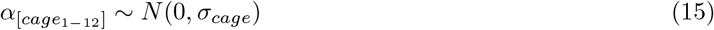

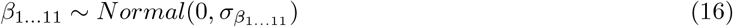

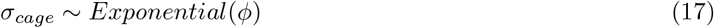

where each *y_i_* is described by a Poisson distribution with mean *λ_i_*. Because the likelihood is not normal, we specify a log link - log(*λ_i_*) - so that the mean can be estimated as a linear function of predictors. This also ensures that the mean will be a positive number, preventing the model from predicting negative spider abundance. In this model, the intercept *α* represents the predicted log mean number of spiders in the treatment with Smallmouth Buffalo on the first sample date. The choice of reference treatment is arbitrary. Choosing Smallmouth Buffalo and the first date as the intercept is the default choice in R (R Core Team 2020) because the treatment is coded first alphabetically (“buffalo”) and first numerically (“2015-06-08”).

#### Priors

As before, we simulated outcomes under three model scenarios, each with different priors (Table 1; Figure 3a-c). Another complication in this model is the log-link, which changes the biological interpretation of the priors. With a log-link the individual model parameters are less intuitive than they would be under a normal likelihood (Bedrick et al. 1996). Under a normal likelihood, a 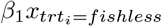 value of 1.5 would indicate that the fishless treatment on 2020-06-08 contains 1.5 more spiders on average than the Smallmouth Buffalo treatment on the same date. With a Poisson likelihood and log-link, the same value first needs to be exponentiated *exp*(1.5) = 4.5 and then interpreted as a multiplier. Thus, a value of 1.5 for the parameter indicates that the fishless treatment contains 4.5 *times* more spiders than the Smallmouth Buffalo treatment on the first sample date. A value of 10 results in 22,026 times more spiders. This is an example of the principle that the prior can only be understood in the context of the likelihood (Gelman et al. 2017).

**Figure 3:**
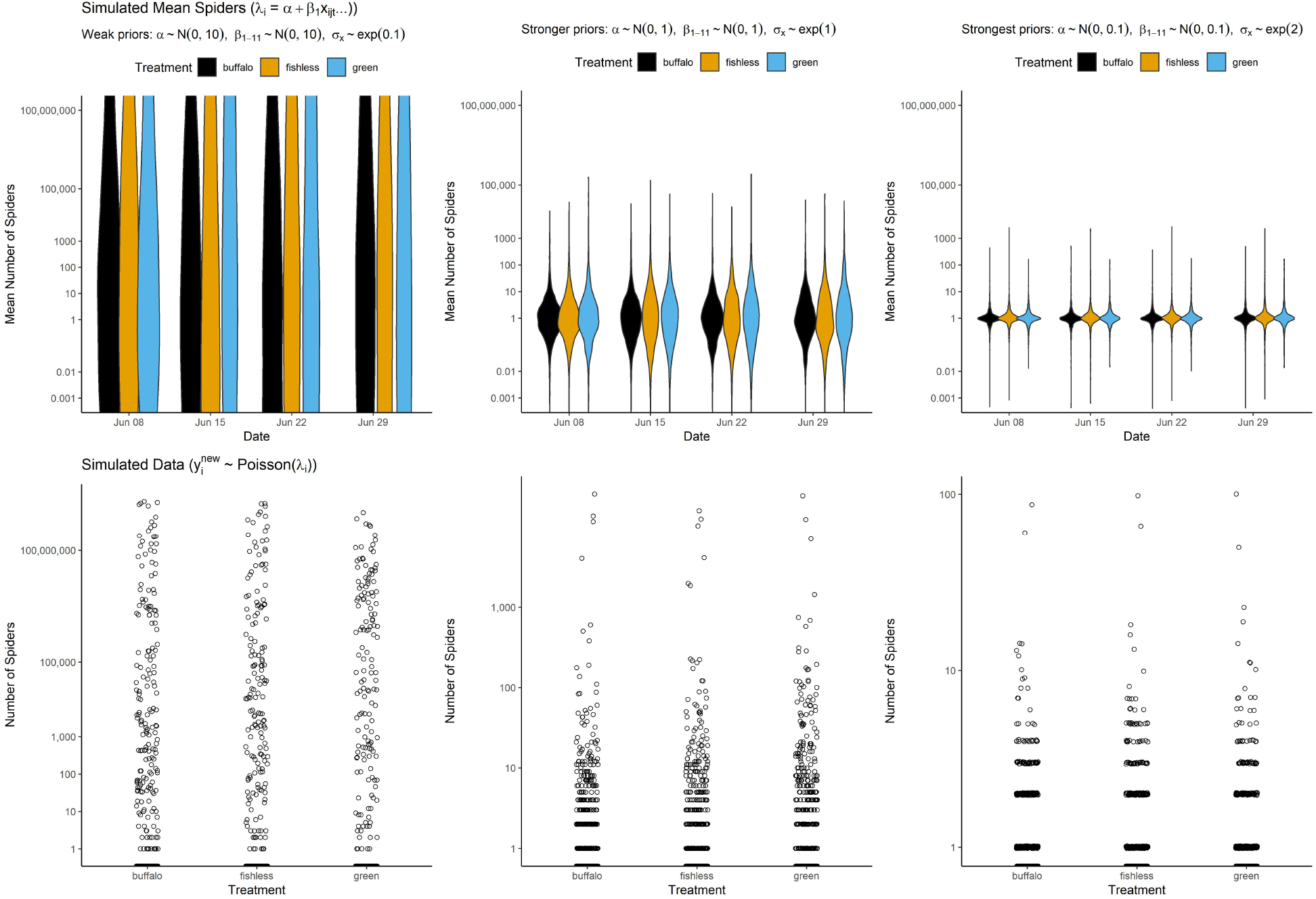
Prior predictive simulations showing the implications of the priors on spider densities above mesocosm cages. Top row: Prior predictive distribution of the number of the mean number of spiders above treatments with either Smallmouth Buffalo, no fish, or Green Sunfish. a) wide priors (*σ_α/β_* = 10, *ϕ* = 0.1), b) stronger priors (*σ_α/β_* = 1, *ϕ* = 1, or c) the strongest priors (*σ_α/β_* = 0.1, *ϕ* = 2). Bottom row: 500 simulations from the prior predictive distribution of the total number of spiders expected for a new cage. Simulations come from the same priors as described above as d) wide priors, e) stronger priors, and f) the strongest priors. To improve visualation, the y-axis for a) is clipped at 0.001 and 1e9.

#### Results

If all we knew was that spiders were counted above 2.32 m^2^ cages but we did not know anything else about the experiment (i.e. the ecosystem, the question, the spider taxa), then we could still use the prior predictive distribution to select more informative priors. The weakest priors place substantial probabilities on values of >100,000 spiders per cage *on average* (Figure 3a), and include a small number of predictions on the final sample date with more than 100 million spiders (Figure 3c). We looked up the range of spider masses (~0.0005 to 170 grams). If we assume our spiders are relatively small, say 0.01 grams, then 100 million spiders would equal 30 tons of spiders. This is approximately equal to the mass of ~6 adult hippopotamus’s (each ~4 tons).

However, in this case we do have valuable prior information. In a previous study using the same cages in the same backwater, (Warmbold 2016) counted between 0 and 2 spiders per cage. The present experiment had a slightly different design, in which a small rope was added to the center of each cage to increase the area of attachment (Warmbold and Wesner 2018). If we assume that the rope will double the number of spiders that could colonize, then it seems reasonable to expect ~ 4 spiders per cage. There is obvious error associated with this, since the experiment was conducted in a different year and a different month. For that reason, we chose the moderate prior (Figure 3b,d) to use in the final model. It places most of the prior probability on values between ~1 to 100 spiders, but also allows for some extreme possibilities of >1000 spiders per cage (Figure 3d). The strongest priors also appear reasonable, placing most of the prior probability between ~1 to 10 spiders, while allowing for up to ~100 spiders in extreme cases (Figure 3c,e).

Figure 3 shows the results after fitting the model to data. Spider counts ranged from 0 to 5 spiders per cage (Figure 4a), resulting in mean spider densities of ~1 to 4 spiders among the date x treatment combinations (Figure 4a). Simulating from the prior and posterior predictive distributions shows the model predictions for the number of spiders we might expect at a new cage (i.e. a cage sampled from this site at another time). Before seeing the data, the model suggested reasonable probabilities of collecting 10 to >100 spiders. After seeing the data, the model suggests that finding ~10 or more spiders would be surprising (Figure 4b).

**Figure 4:**
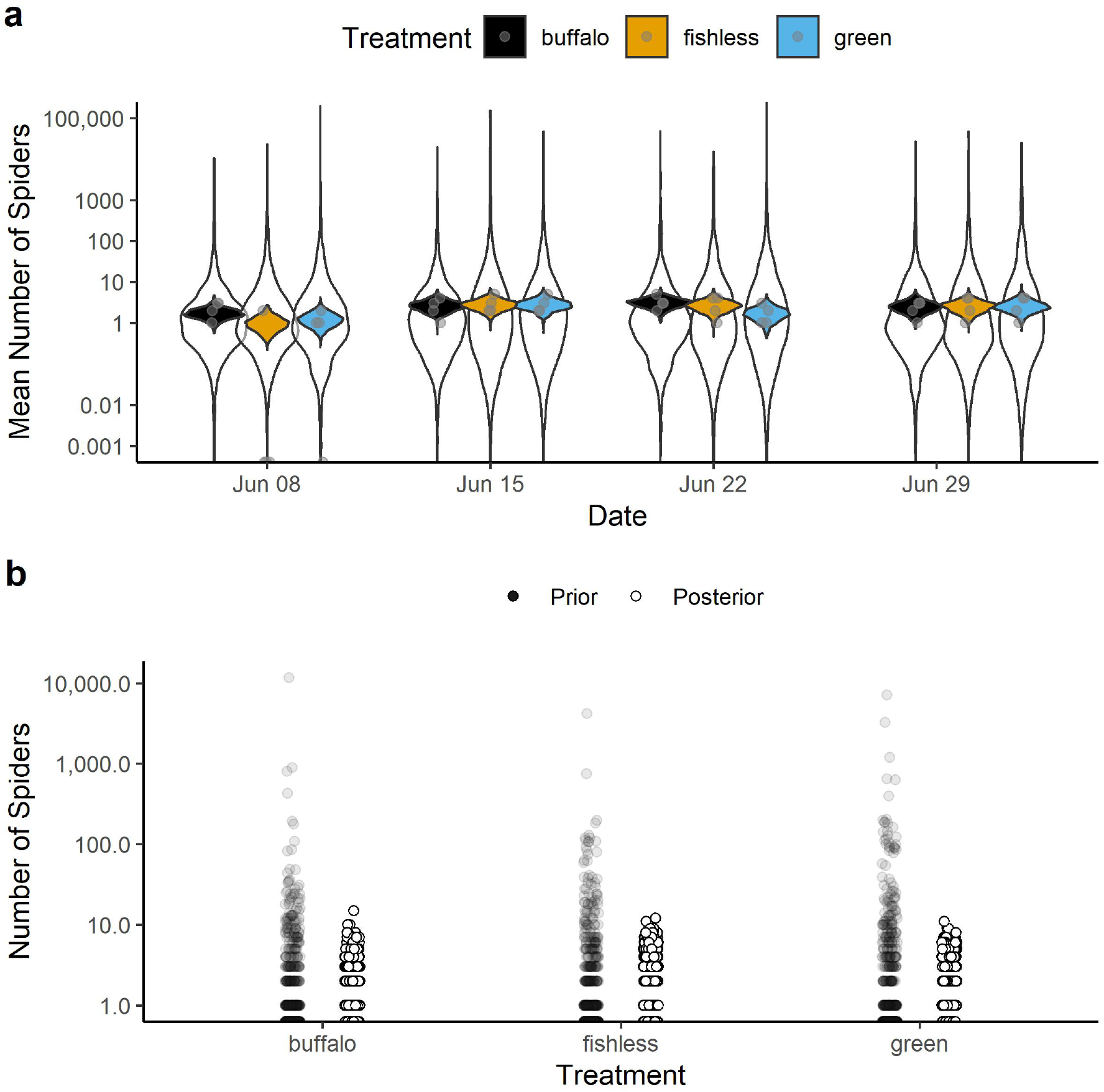
Comparison of the prior and posterior distributions for a) mean number of spiders and b) the conditional prediction of the number of spiders predicted for a new cage from each date x treatment combination. Each violin plot in (a) shows either the prior (white) or posterior (color) distribution with dots as raw data. Each dot in (b) is a simulation (n = 500) of the total number of spiders predicted for a single new cage in each date x treatment combination. The prior is taken from the strong prior in Figure 4b. It is clear that a large amount was learned from the data, as evidenced by the difference between the prior and posterior distributions in both panels.

In addition to the computational and logical benefits of stronger priors as mentioned above, the stronger prior specifications in this model have a clear influence on the posterior (Figure S1). In particular, the stronger prior used in the model is more conservative, pulling the posterior means away from extreme high or low values. As such it acts to prevent overconfidence in large or small effect sizes (e.g., Type M errors) (Lemoine 2019). This skepticism of stronger priors is a benefit that is most apparent with small sample sizes, which are common in ecological studies.

#### Caveats

Each of the 11 *β*’s was assigned an independent prior. An alternative approach would be to assign *β* priors from a multivariate normal distribution (Hobbs and Hooten 2015). In addition, the likelihood assumes that the variance is equal to the mean. An alternative likelihood, such as a negative binomial, would allow us to model variances independently. Finally, the strongest priors we specified overwhelmed the small data set, pulling all treatments towards the same mean, regardless of the data (Figure S1). Whether that is a problem or not depends on how skeptical we are that the cages or treatments would have different numbers of spiders.

## Discussion

Bayesian statistics is increasingly used by ecologists (Ellison 2004, McCarthy and Masters 2005, Hooten and Hobbs 2015, Touchon and McCoy 2016), yet the preponderance of studies continue to rely on diffuse and/or default priors (Lemoine 2019, Banner et al. 2020). Using two case studies with a linear regression and a generalized linear mixed model - two common types of models in ecology (Touchon and McCoy 2016) - we demonstrated how visualization on the scale of the outcome can improve our choices of priors on individual parameters in a Bayesian analysis. From our own experience teaching Bayesian statistics to graduate students (JSW) and the experiences of others (James et al. 2010, Gabry et al. 2019), we suspect that this approach will help to remove confusion or anxiety over choosing more informative priors by aligning the choices more closely to the domain expertise of the users (Bedrick et al. 1996, James et al. 2010).

Choosing priors based on their implications on the outcome scale is not new. Kadane et al. (1980) described a similar approach with normal linear regressions to elicit prior information from experts, and (Bedrick et al. 1996) expanded it to generalized linear models. More recently, (Gabry et al. 2019) used it in a model with random effects to measure global air quality. (Kennedy et al. 2019) used a similar approach for models in cognitive science. A primary difference between the earlier and later uses of prior predictive simulation is the improvement in visualization techniques (Gabry et al. 2019), which makes it easier evaluate prior choices on a visual *distribution* of outcome measures, rather than only point estimates.

Assessing and visualizing priors on the outcome scale of a model makes clear what many current Bayesian approaches emphasize: it is almost never the case that we have absolutely zero prior information (Hobbs and Hooten 2015, Lemoine 2019, Banner et al. 2020). For example, it does not take expertise in ecology or in predator-prey interactions to know that predators cannot eat prey larger than earth, yet this type of impossible prior belief is exactly what many Bayesian models encode with non-informative priors. It *does* take ecological expertise to know whether it is more probable for predators to eat prey that are 2 times larger or 2 times smaller, or whether the log-linear model should have a different functional form (e.g., non-linear). Critiquing priors in this way would, we argue, lead to a much better use of Bayesian methods than current practices that focus on finding the least informative prior (Lemoine 2019, Banner et al. 2020). Even for models with more abstract outcomes than body size (e.g., gene methylation, stoichiometric ratios, pupation rates of a new insect species), it is almost always the case that ecologists have some sense of what reasonable measures might be. After all, it would be impossible to do any sort of study without first knowing what we will measure. Prior expectations of those measures come either from prior experience, the literature, or most often, both.

Visualizing simulations from the prior predictive distribution represents one aspect of the overall Bayesian modeling workflow (Kennedy et al. 2019, Gelman et al. 2020, Schad et al. 2020, Gabry et al. 2019). Like any approach to data analysis, the Bayesian workflow involves iteratively checking assumptions and implications of our model, from data collection and model design to prior choices and model inference (Hooten and Hobbs 2015, Gelman et al. 2020). Traditionally, the role of priors in this workflow has focused on choosing the least informative priors possible, leading to a large body of theoretical and applied literature on development of non-informative priors, such as Jeffrey’s, Horseshoe, or flat priors (Hobbs and Hooten 2015). When prior criticism is used, it is usually done after the model is fit with prior sensitivity analyses and/or plots of prior versus posterior parameters (Korner-Nievergelt et al. 2015). The approach we demonstrate does not obviate the need for these techniques in any sense. Rather, it adopts the approaches that are generally reserved for exploring the implications of the posterior distribution and applies them to the prior distribution. In doing so, it helps to lessen the impact of poor prior distributions later in the analysis workflow.

In ecology, the most closely related application of the approach we describe is for eliciting prior information from a panel of experts (James et al. 2010). However, external elicitation is not practical for most ecological studies, because the data analyst is often also the domain expert (Ellison and Dennis 2010). In other words, most statistical analysis in ecology is done by people (such as us) that are trained in disciplines other than statistics (Touchon and McCoy 2016). As a result, Bayesian analysis in ecology has traditionally been limited to ecologists with advanced statistical and computing capabilities. This is in part because Bayesian analysis is not included by default in popular statistical software, such as R (R Core Team 2020), and also because of the large computing time needed to run Bayesian models relative to frequentist or maximum likelihood approaches. Yet recent improvements in both the MCMC algorithms (Gelman et al. 2015) and the packages used to fit models appear likely to continue the trend of ecologists using Bayesian statistics. For example, with the *brms* package in R (Burkner 2017), this frequentist linear regression - `lm(y \~ x, data = data)` - becomes this Bayesian regression by changing two letters - `brm(y \~ x, data = data)`. This represents the simplest of cases (priors can and should be specified in the `brm()` model), but demonstrates the ease with which fitting Bayesian models is now possible.

An added benefit to choosing more informative priors is that it reduces the computational time needed to fit models, because it limits the parameter space that an MCMC algorithm needs to explore. In the relatively simple models we used here, the computational improvements are likely minimal. But ecologists are using increasingly sophisticated models (Touchon and McCoy 2016), for which the improvements in computational efficiency are likely to be important. An irony in this improvement is that it contradicts a common justification of using non-informative priors to “let the data speak for themselves”. In a model with such priors, much of the “speaking” is done by the priors in the sense of sampling parameter spaces that are incompatible with reasonable data. More importantly, as shown by the first analysis here and by (Gabry et al. 2019), non-informative priors on parameters can become informative for quantities of interest (e.g., average prey sizes that are larger than earth). To rearrange the statement, data can only speak for themselves if the microphone is properly tuned.

## Acknowledgements

This material is based upon work supported by the National Science Foundation under Grant No. 1837233. JSW thanks the students in his graduate Bayesian class for asking challenging questions. He hopes this manuscript provides a long overdue answer. We thank (Brose et al. 2006) for making their data publicly available.

## Supplementary Information

**Figure S1.**
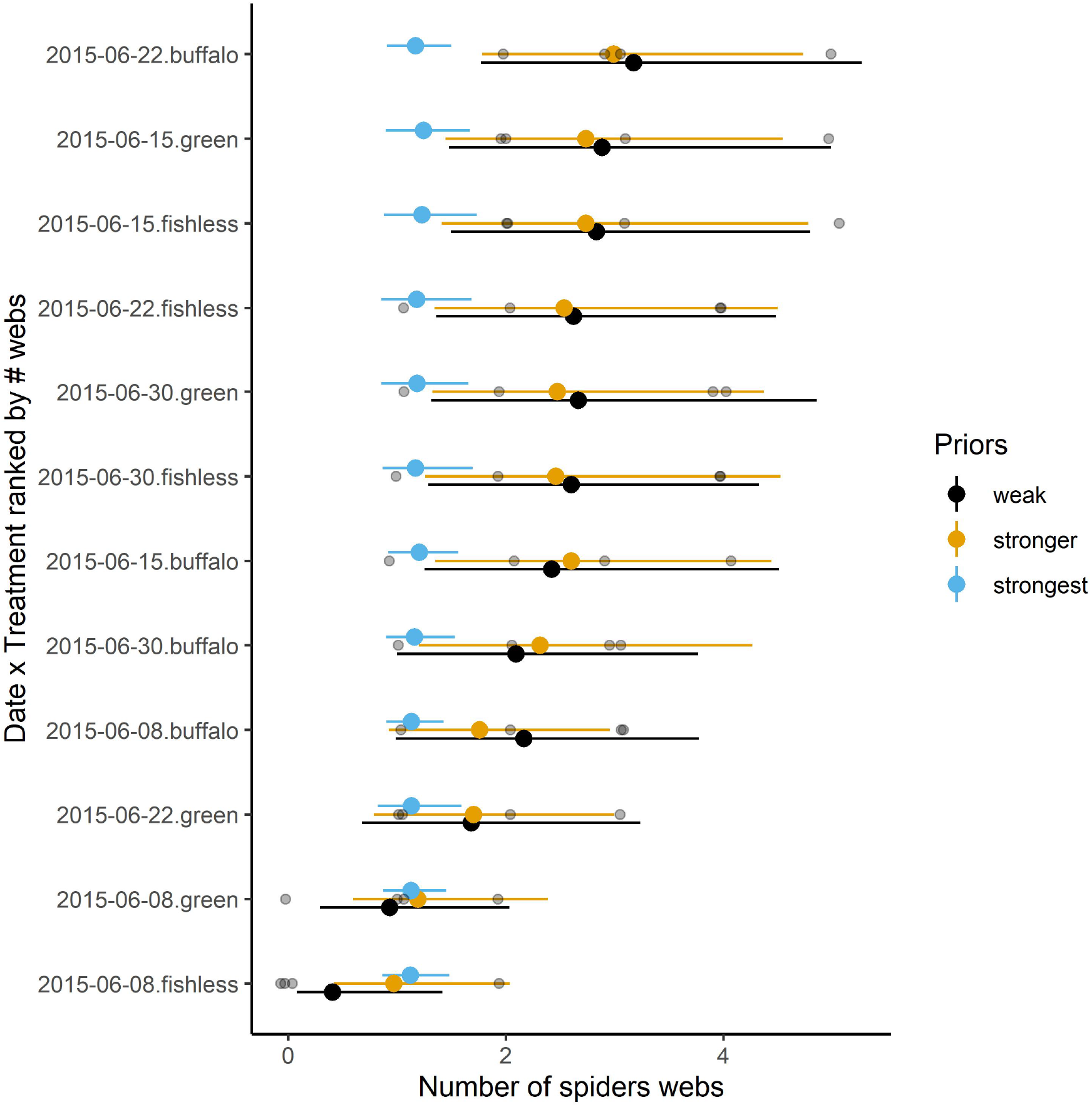
The influence of the prior distributions. Because of the small sample size (n = 4 replicates), the prior specifications affect the posterior. Compared to the weakest prior, the stronger prior is more conservative, pulling each mean towards the prior mean. The strongest prior (blue) essentially overcomes any information in the data. Gray dots are raw data.

